# Reh1 is Required for Nonfunctional 25S RNA Decay

**DOI:** 10.64898/2026.05.05.723076

**Authors:** Caroline Wang, Sham Sunder, Arlen W Johnson

**Affiliations:** Department of Molecular Biosciences, The University of Texas at Austin; Austin, Texas, USA

**Author notes:** corresponding author, 2506 Speedway, NMS 1.246 Department of Molecular Biosciences The University of Texas at Austin Austin, TX 78712-1095.

**Keywords:** REH1, Nonfunctional RNA Decay (NRD), Ribosome Degradation

## Abstract

25S nonfunctional RNA decay (NRD) eliminates 60S ribosomal subunits carrying inactivating mutations in the RNA. However, how cells identify defective subunits has not been described. We recently showed that the zinc-finger protein Reh1 is the last assembly factor to be released from a nascent 60S subunit. We now show that in yeast Reh1 is required for the degradation of 25S NRD substrates. 25S rRNAs carrying mutations in the catalytic center, A2820G or U2954A (A2451 and U2585, respectively in *E coli* numbering), are unstable in wildtype cells but are fully stabilized when *REH1* is deleted. However, not all 25S rRNA mutations are recognized by Reh1. Ribosomes with a truncated L1 stalk engage in translation but cannot support viability. These ribosomes display a half-life indistinguishable from wild-type rRNA, suggesting that yeast does not have a robust surveillance system for such mutant ribosomes. Deletion of *REH1* also has no impact on the levels of defective 18S rRNA. These results indicate that Reh1 and 25S NRD are specific for mutations in or near the catalytic center of the ribosome.

## Introduction

The faithful expression of a cell’s genome depends on accurate translation by the ribosome. The ribosome itself is an intricate assemblage of RNA and protein which carries out two distinct functions: the small subunit (40S in eukaryotes) decodes mRNA while the large subunit (60S) synthesizes polypeptides. The process of translation is highly dynamic, requiring the rapid interrogation and discrimination of tRNAs, and involves small scale rRNA movements within the functional centers as well as large-scale interdomain movements and subunit ratcheting. Consequently, ribosome function depends on the precise synthesis and assembly of its components to support the dynamic process of translation with high accuracy and speed.

In eukaryotes, ribosome subunit assembly is a complex pathway, occurring largely in the nucleolus with additional maturation in the nucleoplasm and subsequently in the cytoplasm after export from the nucleus. Most defects in rRNA processing or assembly in the nucleus are recognized and efficiently eliminated by pathways requiring the nuclear RNA exosome [1–3]. Cells have evolved elaborate systems to ensure the functional integrity of ribosomes. Indeed, even the absence of a rRNA methylation can block progression of the assembly pathway[4]. Nevertheless, some subtle defects, such as inactivating mutations in 18S rRNA in the decoding center of the 40S subunit or in 25S rRNA in the peptidyl transferase center (PTC) of the 60S subunit can evade detection during assembly in the nucleus. Such defective ribosomes can escape to the cytoplasm where they are subjected to surveillance by non-functional ribosomal RNA degradation (NRD) pathways[5,6].

Interestingly, mutations in the RNAs of the small and large subunit are recognized and degraded by separate pathways. Early work from the Moore lab showed that mutations in the decoding center of 18S rRNA are eliminated by a translation-dependent mechanism that involves factors of the No-Go Decay pathway, including Asc1, Dom34 and Hbs1[5,7,8]. One well-characterized 18S NRD substrate is A1775C in 18S rRNA, corresponding to the universally conserved A1492 in E coli, which is important in recognition of cognate tRNAs during translation[9]. In yeast, ribosomes carrying the A1775C mutation arrest at the start codon[10]. Subsequent work has revealed that defective 80S ribosomes are sequentially ubiquitinated on uS3 first by Mag2 and then by Hel2 or Fap1.[10,11] The poly-ubiquitination of uS3 results in the recruitment of the ribosome quality control trigger complex which separates the ribosomal subunits to facilitate 40S degradation. In human cells, 18S NRD operates through the integrated stress response and requires ZNF10, the human counterpart of yeast Mag2[12], indicating conservation of the pathway for eliminating defective 40S subunits.

In contrast to 18S NRD, degradation of 25S rRNA by the 25S NRD pathway does not appear to require ongoing translation, unlike 18S NRD[7]. 25S NRD also does not require No-Go decay factors or the RQT complex[7] which raises the question of how cells detect non-functional 60S subunits if they are not engaged in translation. Two well-characterized 25S NRD substrates are A2820G and U2954A (corresponding to nucleotides A2451 and U2585 in E coli)[5]. Work by the Kitabatake and Ohno groups has shown that degradation of both 25S NRD substrates requires the cullin E3 ligase complex of Rtt101 and Mms1[6]. This complex also contains Hrt2 which tethers the E2 ligase Cdc34 and utilizes Crt10 as an adapter for substrate recognition. The Rtt101-Mms1 complex is also important in DNA repair pathways in which Mms22 is used as the adapter for substrate recognition[13]. Deletion of *MMS1* leads to sensitivity to the DNA alkylating agent methyl-methanesulfonate[14], which also modifies RNAs and can lead to induction of stress pathways through ribosome dysfunction[15]. Ultimately, defective 60S subunits are degraded by the proteasome[16].

Although some 25S NRD substrates, such as U2954A appear to arrest as free 60S subunits and do not appear to engage with 40S subunits to form 80S monosomes, other 25S NRD substrates, such as A2820U, can progress further and engage with 40S subunits. It has been reported that Cdc48-Npl4-Ufd1 complex splits these defective 60S subunits from 40S subunits during 25S NRD[16]. Because the conclusion that translation is not needed for 25S NRD was derived from work using the U2954A mutation, which does not engage with 40S subunits[5], it remains a possibility that mutant 25S that engage with 40S require translation for their degradation. Despite our understanding of the downstream fate of 25S NRD substrates, a major outstanding question has been what factors on the 60S subunit signal that it is defective and in what context is the defective subunit recognized, if not during translation.

During ribosome assembly the insertion of the ribosomal protein (rprotein) uL16 completes the catalytic center of the ribosome[17,18]. An internal loop of uL16, the P site loop, is a dynamic element which interacts with ligands in the A and P sites and is required for efficient translation[19,20]. The accommodation of this loop into its proper position triggers the release of the export adapter Nmd3[17] and sets the stage for a “test drive” in which molecular mimics of translation machinery engage the subunit[21]. During the test drive the small protein Sdo1 (SBDS in humans) binds in the P site and activates the GTPase Efl1, a paralog of the translation translocation factor eEF2, which evicts the antiassociation factor Tif6 (eIF6 in higher eukaryotes)[22,23]. Because Tif6 blocks association with the small subunit, the release of Tif6 “licenses” the subunit for translation. We showed previously that mutations in the P site loop of uL16 block the test drive, implying that cells interrogate the integrity of the P site before licensing the subunit[21].

We have recently shown that in yeast Reh1, a paralog of the assembly factor Rei1, serves as a quality control factor, and is required for eliminating ribosomes containing mutant uL16, in which the P-site loop of uL16 has been deleted[19]. The mutant uL16 inserts into ribosomes during assembly but blocks the test drive and prevents the release of Tif6 and the export factor Nmd3. These mutant ribosomes can be released into translation by mutations in Nmd3 and Tif6 that weaken their affinities for the ribosome. Surprisingly, we found that when these defective ribosomes engage 40S subunits and enter translation, they are relatively stabilized. Thus, in the case of uL16 P-site mutants, defective 60S subunits appear to be surveilled predominantly during the test drive and not during translation. Furthermore, Reh1 is required for the selective removal of 60S subunits containing mutant uL16[19]. In human cells, a recent CRSPR screen for quality control factors using the corresponding uL16 mutation in human cells identified the zinc-finger protein ZNF574[24]. As yeast does not have an obvious counterpart for ZNF574 it is unclear what the functional relationship is between ZNF574 and Reh1, but it is likely that ribosome surveillance pathways in human cells are more complicated than those in yeast.

Here, we have asked if Reh1 in yeast also acts on canonical 25S NRD substrates in addition to mutant uL16-containing subunits. We found that in the absence of Reh1 the 25S NRD substrates A2820G and U2954A become fully stabilized, with half-lives indistinguishable from WT. The 25S mutant subunits accumulate primarily at 60S, suggesting that they are recognized as nascent 60S subunits before they engage in translation, consistent with the previous observation that 25S NRD is independent of ongoing translation. In contrast, deletion of *REH1* had little impact on the 18S NRD substrate A1775C. Deletion of *REH1* also had no impact on a different 25S rRNA mutant lacking the L1 stalk, which is distal to the PTC. Surprisingly, although 25S deleted of the L1 stalk cannot support growth, the mutant rRNA displays a half-life like that of wild-type 25S in WT cells, suggesting that although these mutant 60S enter translation, there is not a robust degradation pathway for the removal of subunits lacking the L1 stalk. We conclude that Reh1 is a dedicated quality control factor for 60S subunits with defects in the catalytic center. We propose that Reh1 is the ribosome-proximal factor that initiates 25S NRD.

## Results

### Establishing 25S NRD substrate vectors

To determine if *REH1* is required for 25S NRD, we first generated 25S NRD substrates modeled on previous work [5]. We introduced the mutations A2820G and U2954A (A2451 and U2585 in E coli numbering, respectively) into a high copy (2 micron) vector that expresses the rDNA locus under control of its native RNA Pol I promoter. We also engineered neutral tags in 18S and 25S to be able to normalize 25S expression to 18S expression from the same vector (Figure 1A). This is possible because 18S and 25S are derived from the same primary transcript to ensure stoichiometric production of the 40S and 60S subunits. We employed an oligo tag in expansion segment (ES) 2 of 18S as previously described (PMC20288) and engineered a new oligo tag, (5’-GAAGAGCCATTGCACTCCGGTTCTTCTGCAG) which included a binding site for human U1A protein, between residues 137 and 138 in ES5 of 25S. Cells expressing rDNA with a single tag in 25S or tags in both 25S and 18S from vectors as their sole copy of rDNA grew similarly to cells expressing untagged vector-borne rDNA (Figure 1B). We then compared expression of wild-type and mutant rRNAs using two color fluorescent northern blotting (Figure 1C).

**Figure 1.**
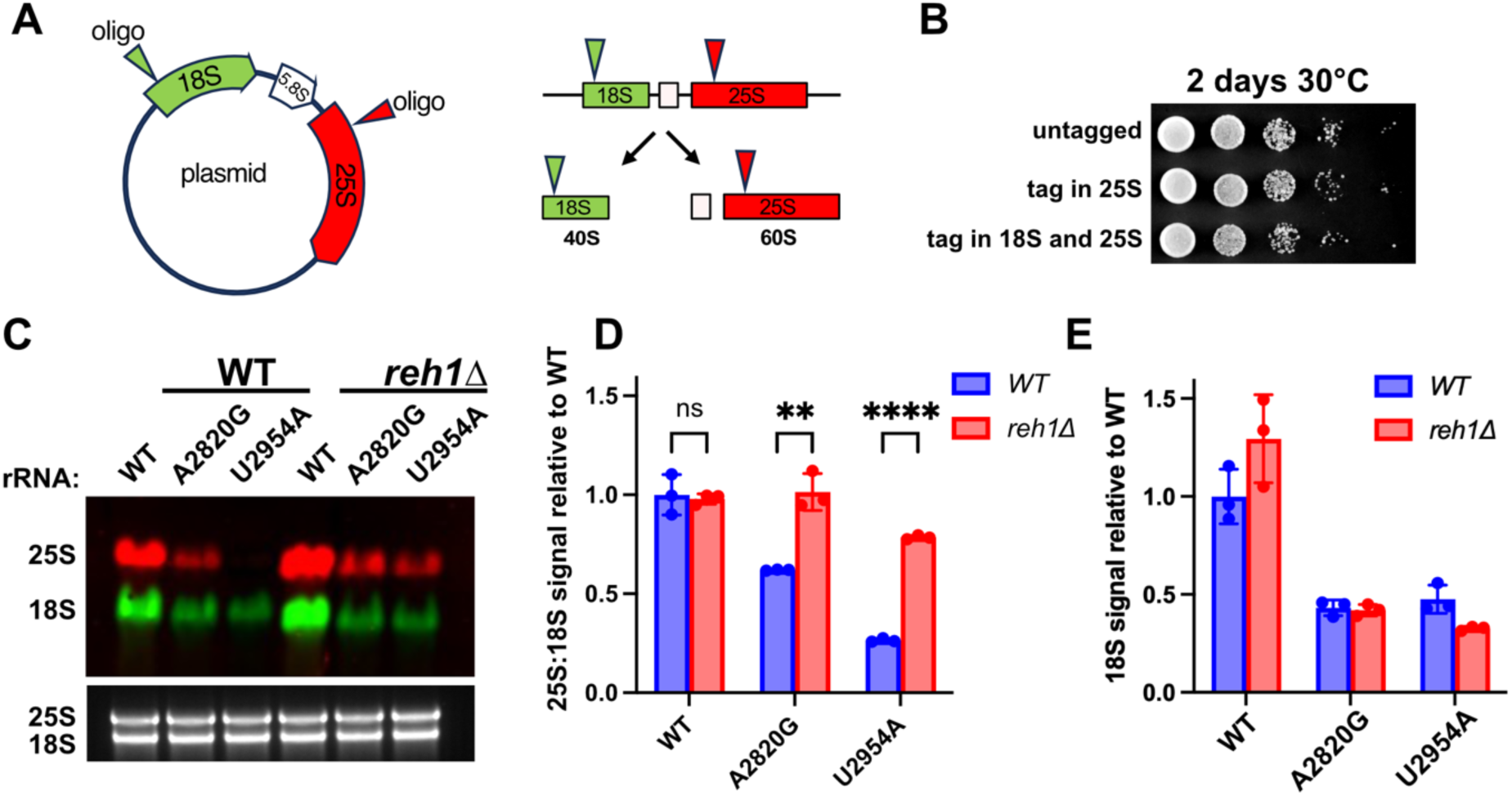
Steady state analysis of 25S NRD substrates. **A.** Cartoon of rDNA vector showing the relative positions of oligonucleotide insertions in 18S and 25S rRNAs for northern blotting. Transcription of the rDNA locus yields a primary transcript that is then processed into 18S (green) and 25S rRNAs (red), the major RNAs of the 40S and 60S subunits, respectively. **B.** Serial dilution of cells deleted for the rDNA locus and containing plasmids expressing ribosomes that are untagged pJD211.LEU2, with 25S tag only (pAJ1181) or containing both 18S and 25S tags (pAJ5659) as the sole source of rRNA. **C.** Representative northern blot of wild-type or *reh1Δ* strains containing 2µ plasmids expressing either WT or A2820G or U2954A 25S probed with fluorescent oligos specific for 25S (red) or 18S (green) rRNA (top). Total RNA from a TAE-agarose gel stained with SYBR^TM^ Safe (bottom). Equal amounts of total RNA were loaded in each lane. **D.** The 25S signal was quantified by calculating the ratio of 25S to 18S fluorescent signals, normalized to the ratio for 25S:18S rRNA in WT cells. RNAs were prepared from 3 experimental replicates. **E.** The 18S signal was quantified by calculating the ratio between 18S from the mutant vector compared to 18S from WT and normalized to WT rRNA in WT cells. Quantitation was from three experimental replicates.

We compared the ratios of 25S to 18S and normalized values to the ratio of the wild-type rRNA which we arbitrarily set to one, assuming that wild-type 25S and 18S were expressed in equal stoichiometry (Figure 1D). Normalizing to rRNA expressed from the same vector accounted for the variable expression from these vectors. We found that the levels of the mutant rRNAs were reduced compared to wild-type with the A2820G mutant present at 69% the level of wild-type and U2954A at 24% of wild-type (Figure 1C,D). These results are qualitatively similar to what has been reported previously for A2820G and U2954A mutant 25S rRNA. However, the magnitude of the effects of the vectors used here was less than what was reported previously, where A2820G and U2954A mutant rRNAs were 20% and 10% of the level of wild-type rRNA, respectively (LaRiviere). Additionally, we noticed that the 18S signal from mutant 25S-borne vectors was significantly lower than the 18S signal from wild type 25S-borne vectors. We found that 18S originating from vectors expressing mutant 25S was expressed at lower levels than 18S from the vector bearing wild-type 25S; the levels of 18S from the A2820G-expressing vector was 43% of that of WT and U2954A was 48% of WT (Figure 1E). We suspect that the differences in expression from wild-type and mutant-expressing vectors results from a fitness defect imposed by the mutants, resulting in reduced copy number of high-copy vectors used in this work (see below).

In an attempt to understand the apparent differences in expression of mutant rRNAs derived from the constructs we generated compared to those used previously[5], we considered that different overall expression levels from the two vector sets could somehow contribute to the differences in expression of the mutant RNAs. For example, if the rRNA was expressed at higher levels from the vectors we made, turnover machinery could be overwhelmed, resulting in the higher apparent steady state levels of mutant rRNA that we found. One significant difference between the vector sets is the use of different promoters. While the vectors generated for this work utilized the native RNA Pol I promoter, previous work relied on expression from the strong constitutive *PGK1* RNA PolII promoter[5]. To compare expression between the plasmid sets, which contained different hybridization tags in 18S and 25S, we moved the 18S tag alone or with the 25S tag from the wild-type *PGK1* vector into the wild-type RNA PolI vector and compared expression levels of 18S. We found that the RNA PolI-dependent vectors expressed 18S at approximately 66% the level of the *PGK1* vectors (Supplemental Figure S1A,B). Because the PGK1-dependent vectors express at higher levels than the PolI vectors, we do not think that the difference in steady state levels of mutant 25S expressed from the PGK1 vector set vs the RNA PolI set can be explained by differences in overall expression.

We then examined expression of mutant 25S rRNA from the original *PGK1* vectors from the Moore lab and found that the A2820G and U2954A mutants expressed from the *PGK1* promoter showed 25S:18S ratios of 0.69 and 0.25, respectively, which was essentially identical to the values measured for RNA PolI vectors of 0.69 and 0.24 (Supplemental Figure S2 A,B). Tellingly, when normalized to *SCR1* RNA rather than 18S rRNA, as was done in the previous work, the apparent expression of A2820G and U2954A levels dropped to 40% and 17% of wild-type levels, respectively, approaching what was reported previously (supplemental Figure S2C). We also examined the overall expression from the *PGK1* vectors by comparing 18S levels and found that vectors expressing mutant 25S rRNA also showed overall reduced expression compared to wild-type as we observed for the RNA PolI vectors (Supplemental Figure S2D). We conclude that the apparent differences in mutant 25S levels measured with the two vector sets is a result of different methods or normalization and that normalizing to 18S expressed from the same vector-borne transcript is critical to account for the variable expression of these mutants.

### Reh1 controls the levels of 25S NRD substrates

We recently identified the pre-60S-associated protein Reh1 as a quality control factor that is required for elimination of ribosomes containing a mutant uL16 protein lacking the P site loop[19]. This loop of uL16 approaches the catalytic center and interacts with ligands in the P and A sites [19,20]. Whether or not the activity of Reh1 is limited to uL16 mutants or acts on a broader range of 60S mutants remained an open question. To this end, we asked if Reh1 promotes the elimination of 60S subunits carrying inactivating mutations in the rRNA. Indeed, we found that in the absence of Reh1, the ratio of 25S to 18S rRNA rose from 0.69 to the wild-type ratio of 1.0 for the A2820G mutant and from 0.27 to 0.79 for the U2954A mutant (Figure 1C, D). Importantly, when normalized to SCR1, the restoration of 25S:18S ratios was not evident because this analysis does not account for the reduced expression from these vectors (Figure S2C). Thus, Reh1 is required for the reduced expression of ribosomes carrying mutations in rRNA, in addition to mutations in uL16.

Surprisingly, although the ratio of 25S to 18S was largely restored, the overall expression from the mutant-expressing vectors in the *reh1*Δ strain, as measured by 18S levels, remained low (Figure 1E). It seems likely that rescuing 60S from degradation may have negative consequences for the cells, imposing an additional fitness burden that suppresses expression of the mutant vectors. To test if the mutant rRNAs confer a fitness defect on cells, we expressed them under control of a galactose inducible *GAL7* promoter in wild-type and *reh1Δ* cells. Indeed, the A2820G and U2954A mutants conferred strong growth defects in *reh1*Δ cells but not in WT cells where they appeared to show no or a very modest impact on growth (Supplemental Figure S3). This result is consistent with the idea that despite the restoration of 25S:18S ratio in a *reh1*Δ mutant, the mutant rRNAs impose a fitness burden.

### Reh1 does not monitor defective 60S lacking the L1 stalk or 40S with mutant 18S rRNA

We next asked if the ability of Reh1 to sense defective rRNA encompasses mutations in other areas of the ribosome. The uL16 mutation we described previously as well as A2820G and U2954A rRNA mutations are all close to or within the catalytic center of the 60S subunit. For comparison, we chose an additional 25S mutant, which produces a 60S subunit lacking the L1 stalk (ΔL1 stalk) [25]. The L1 stalk is distal to the catalytic center but important for interaction with ligands in the E site, including tRNAs and the elongation factor eIF5A[20,26,27]. We previously showed that ribosomes lacking the L1 stalk can enter translation but cannot support growth[25]. To test if Reh1 is required for 18S NRD, we also generated a mutation in the decoding site of the small subunit 18S rRNA, A1775C (A1492 in E coli numbering). This mutant has previously been shown to cause rapid turnover of 18S rRNA in yeast[5] but is detected by a pathway distinct from that which deals with defective 25S rRNA[5,7,8]. Both mutants were expressed at low levels relative to wild-type rRNAs in a wild-type strain background (Figure 2A). The reduced expression of the ΔL1 stalk mutant resulted from reduced expression of the mutant 25S relative to 18S (Figure 2B) as well as reduced total expression from the vector, as seen in the reduced 18S signal (Figure 2C). For the A1775C mutant, we observed reduced expression of mutant 18S relative to 25S expressed from the vector (Figure 2D) as well as an overall reduction in expression from the vector, as seen by the reduced level of 25S from the mutant versus wild-type vector in wild-type cells (Figure 2E). Importantly, deletion of *REH1* did not result in a significant increase in the 25S:18S ratio for the ΔL1 stalk mutant. There was slight but statistically significant increase in the 18S:25S ratio for the 18S A1775C rRNA mutant. However, the increase was small compared to that observed for 25S NRD substrates and seems unlikely to result from the direct activity of Reh1. These results demonstrate that Reh1 does not act broadly on any mutant ribosome but appears to be specific defects in the functional center of the large subunit, regardless of whether they are protein or RNA mutations.

**Figure 2.**
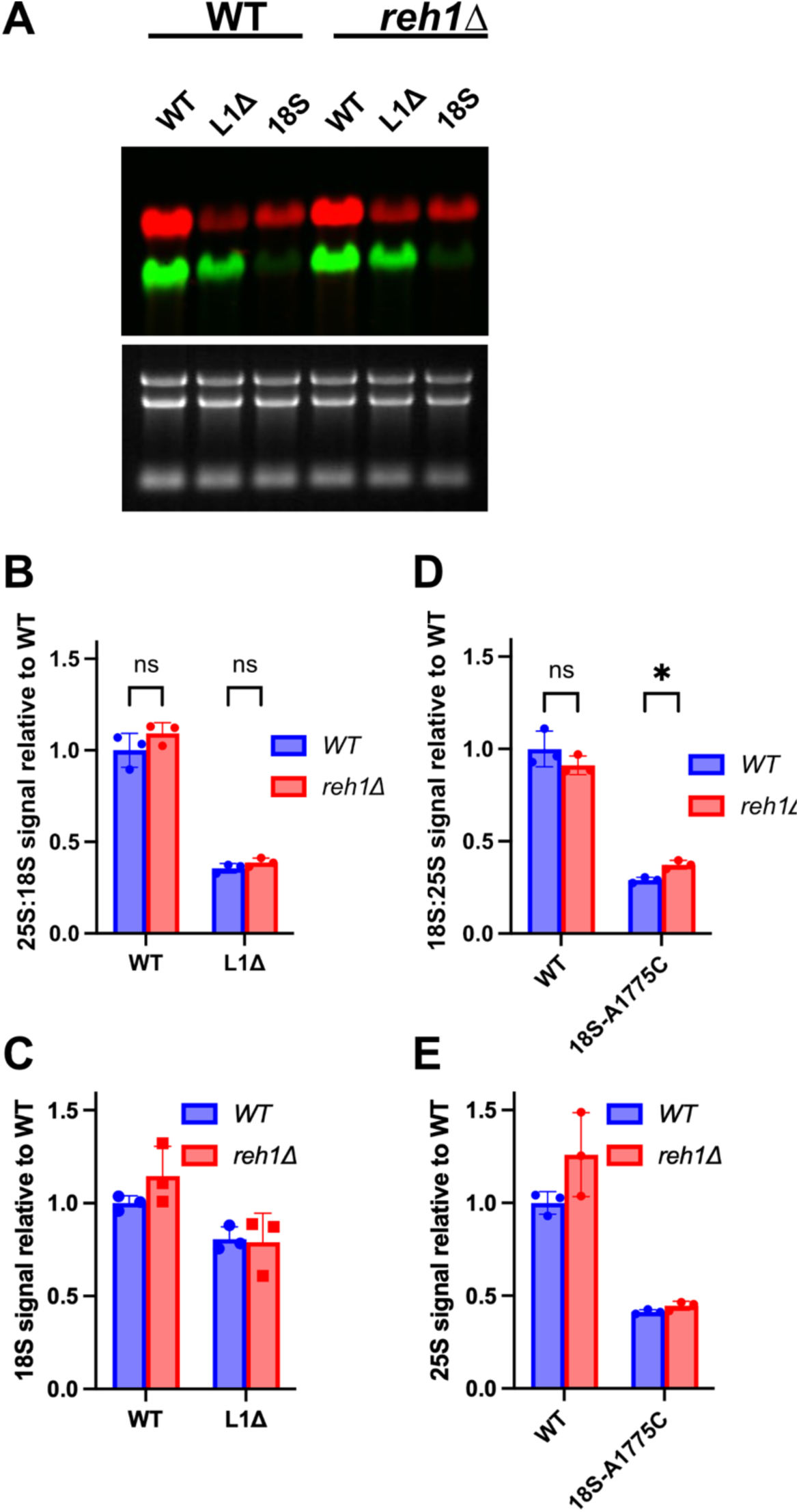
Reh1 is unable to detect defects outside the catalytic center. **A.** Representative northern blot assay of RNA from wild-type or *reh1Δ* strains containing 2µ plasmids expressing either WT or mutated L1-stalk (labeled as L1Δ) 25S rDNA or WT or A1775C 18S rDNA (labeled as 18S). Bottom panel: SYBR^TM^ safe-stained gel of total rRNA. **B.** Quantification of the ratios of 25S to 18S signals in strains containing vector-borne WT or L1Δ rRNA, normalized to WT 25S to 18S ratio in WT cells. RNAs were prepared from 3 experimental replicates. **C.** The ratios of 18S signal in WT and *reh1Δ* cells is plotted relative to the signal of WT 18S rRNA in WT cells. **D.** Quantification of the ratios of 18S to 25S signals in strains containing vector-borne WT or A1775C rRNA, normalized to WT 18S to 25S ratio in WT cells. **C.** The ratios of 25S signal in WT and *reh1Δ* cells is plotted relative to the signal of WT 25S rRNA in WT cells. Quantitation was from three experimental replicates.

### The Reh1 pathway can be saturated

We considered that high levels of expression from high copy vectors may saturate a quality control pathway, reducing the difference in levels of mutant rRNAs compared to WT. To test this idea, we expressed wild-type and mutant rRNAs from low copy centromeric vectors in wild-type and *reh1*Δ cells (Figure S4). Expression from low copy vectors significantly reduced the levels of mutant rRNAs compared to wild-type, as anticipated. Whereas A2820G and U2954A were expressed at 69% and 27% of the levels of wild-type when expressed from 2 micron vectors, those levels fell to 22% and 16% when expressed from centromeric vectors (Figure S4B). This result suggests that elevated levels of expression overwhelm the quality control pathway. In *reh1*Δ cells, the ratios of 25S to 18S returned nearly to WT rRNA-levels, similar to what we observed for 2 micron vectors. We also did not see reduced expression of 18S from the mutant centromeric vectors (Figure S4C), unlike what we observed for the 2 micron series of vectors. However, deletion of Reh1 had a profound effect on overall expression from the entire set of centromeric vectors (Figure S4C), with wild-type rRNA being expressed more than 3-fold higher in *reh1*Δ compared to wild-type cells. The mutant vectors also showed similarly high levels of expression. These observations suggest that expression of the rDNA vectors in the nucleus is being modulated by feedback reliant on Reh1 in the cytoplasm. We are currently working to understand the mechanism of this feedback.

### Reh1 is required for the rapid decay of some NRD substrates

So far, we have only examined steady state levels of rRNAs which is the net result of transcription and degradation and does not explicitly address whether or not the mutant rRNAs display altered stability or if Reh1 plays a role in their stability. Consequently, we measured the half-lives of wild-type and mutant RNAs in wild-type and *reh1*Δ cells. To do this, we used vectors expressing rDNA under control of the *GAL7* promoter. Cells carrying these vectors were cultured in the presence of galactose to induce expression and then shifted to glucose-containing medium to repress *GAL7*-dependent transcription. RNA was collected starting at the shift to glucose (0min) and then at 30, 60, and 120 minutes post-shift to measure degradation of the vector-borne RNAs.

Because we loaded constant amounts of total RNA, the RNAs that were expressed prior to time zero decrease both due to decay over time and also due to dilution with newly synthesized ribosomes. To circumvent the issue imposed by dilution, we plotted half-lives of 25S relative to 18S, normalizing values to one at time zero (Figure 3A). This measures relative half-lives rather than absolute half-lives. However, because wild-type ribosomes have a half-life of several hours in yeast (Supplemental Figure S5), 18S levels serve as a proxy for dilution. A similar strategy of normalizing to rRNA expressed from the same vector was reported previously[8]. The A2820G mutant, which displayed an intermediate steady state level, displayed a modestly reduced relative half-life of 188 minutes and this mutant rRNA was fully stabilized by deletion of *REH1* (Figure 3B). The U2954A mutant, whose steady state level was lowest in wild-type cells among the mutants tested showed a dramatically shortened relative half-life of 25 minutes (Figure 3C). Remarkably, the U2954A mutant 25S was fully stabilized in *reh1*Δ cells.

**Figure 3.**
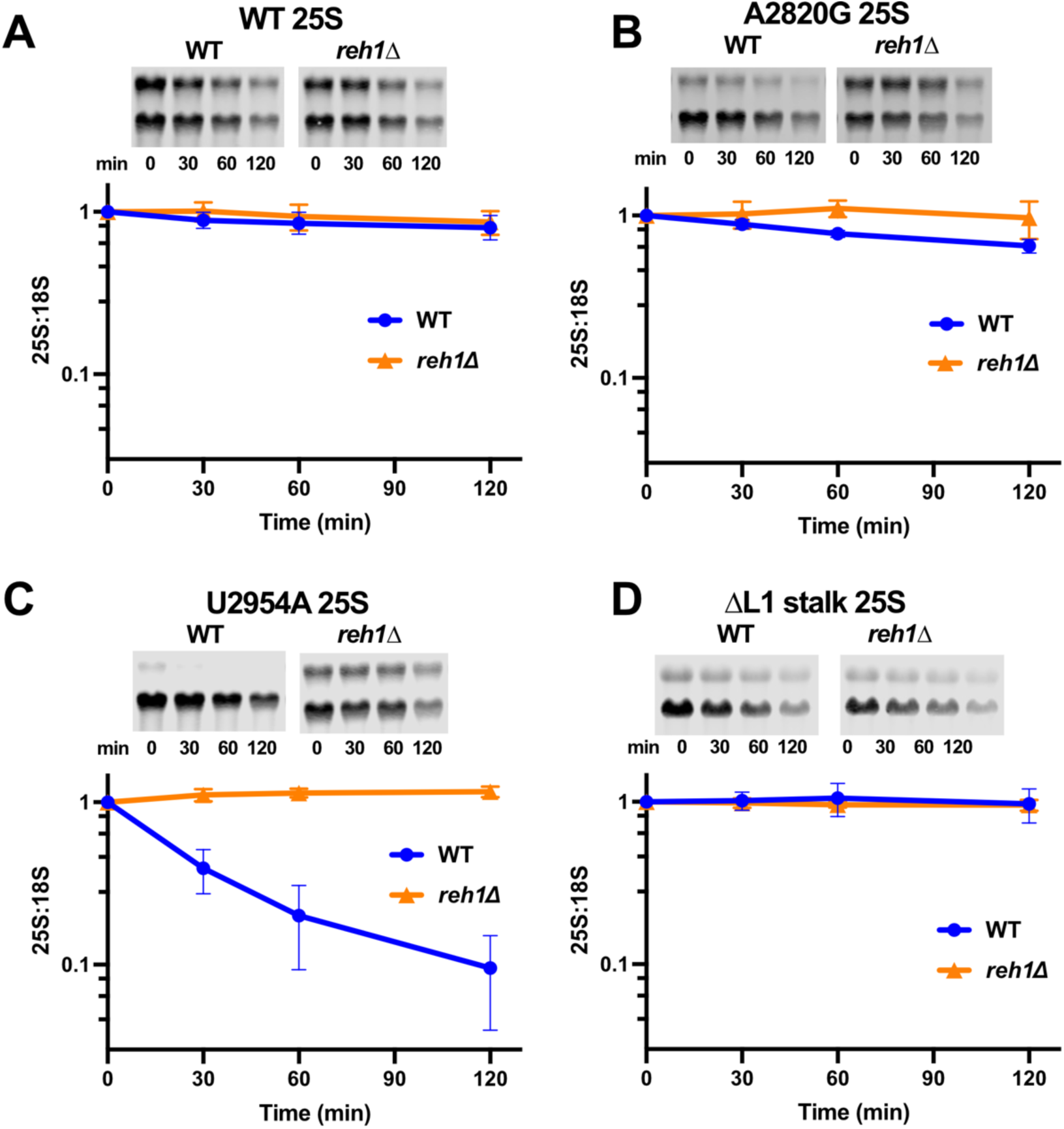
Time course analysis of rRNA stability. WT or mutant rRNAs were expressed from high copy vectors under control of the *GAL7* promoter in WT or *reh1Δ* cells. Cells were harvested at the indicated times after repression of expression by the addition of glucose and RNA was prepared and analyzed by northern blotting. Equal amounts of total RNA were loaded in each lane. Representative blots are shown above each panel and quantification from three experimental replicates is plotted. **A.** WT rRNA, **B.** A2820G, **C.** U2954A and **D.** ΔL1 stalk. Quantitation was from three experimental replicates.

Interestingly, the ΔL1 stalk mutant, which showed low steady state levels of RNA, did not show a statistically significantly decreased relative half-life in wild-type cells and was unaffected by deletion of *REH1* (Figure 3D). The low steady state levels of the ΔL1 stalk mutant are likely due to its reduced rate of nuclear export[25], and nuclear degradation. This result suggests that yeast cells do not have an efficient mechanism for removing these ΔL1 stalk mutant ribosomes despite their inability to support viability. The complete stabilization of the U2954A and A2820G mutants suggests that degradation of the mutants RNAs is entirely dependent on Reh1 and that there is not a redundant pathway for degradation of these ribosomes.

### 25S NRD substrates accumulate at 60S in the absence of Reh1

We have recently shown that uL16 mutant ribosomes accumulate at 60S and weakly enter translation in the absence of Reh1. To determine if Reh1 also gates the release of ribosomes carrying nonfunctional rRNA mutations, we monitored the mutant subunits in sucrose gradients. Previous work from the Moore lab showed that A2820G mutant ribosomes mildly accumulate at 60S but also enter polysomes whereas the U2954A mutant is strongly arrested at 60S[5]. To monitor mutant ribosomes, we used sucrose gradient sedimentation analysis and northern blotting to detect vector-borne 25S and 18S rRNAs. Wild-type 25S was observed at 60S, 80S and polysomes in wild-type and *reh1*Δ cells (Figure 4B). The A2820G mutant was also observed in 60S, 80S and polysomes in wild-type cells, but in comparison to wild-type rRNA, the polysome signal was reduced (Figure 4B). This sedimentation was similar to what was reported previously for this mutant. The distribution of A2820G mutant rRNA was similar in WT and *reh1*Δ cells with the exception that it was slightly increased in the 60S fraction (Fig 4B, fraction 5). The distribution of U2954 mutant rRNA was also similar between wild-type and *reh1*Δ cells with the exception of an increase in the 60S fraction (Figure 4C, fraction 5). Lastly, the ΔL1 stalk mutant showed a similar distribution in wild-type and *reh1*Δ cells (Figure 4D), consistent with our finding that deletion of *REH1* has no impact on the levels of this mutant. Together, these results show that Reh1 primarily affects the levels of mutant rRNA in the free 60S fraction.

**Figure 4.**
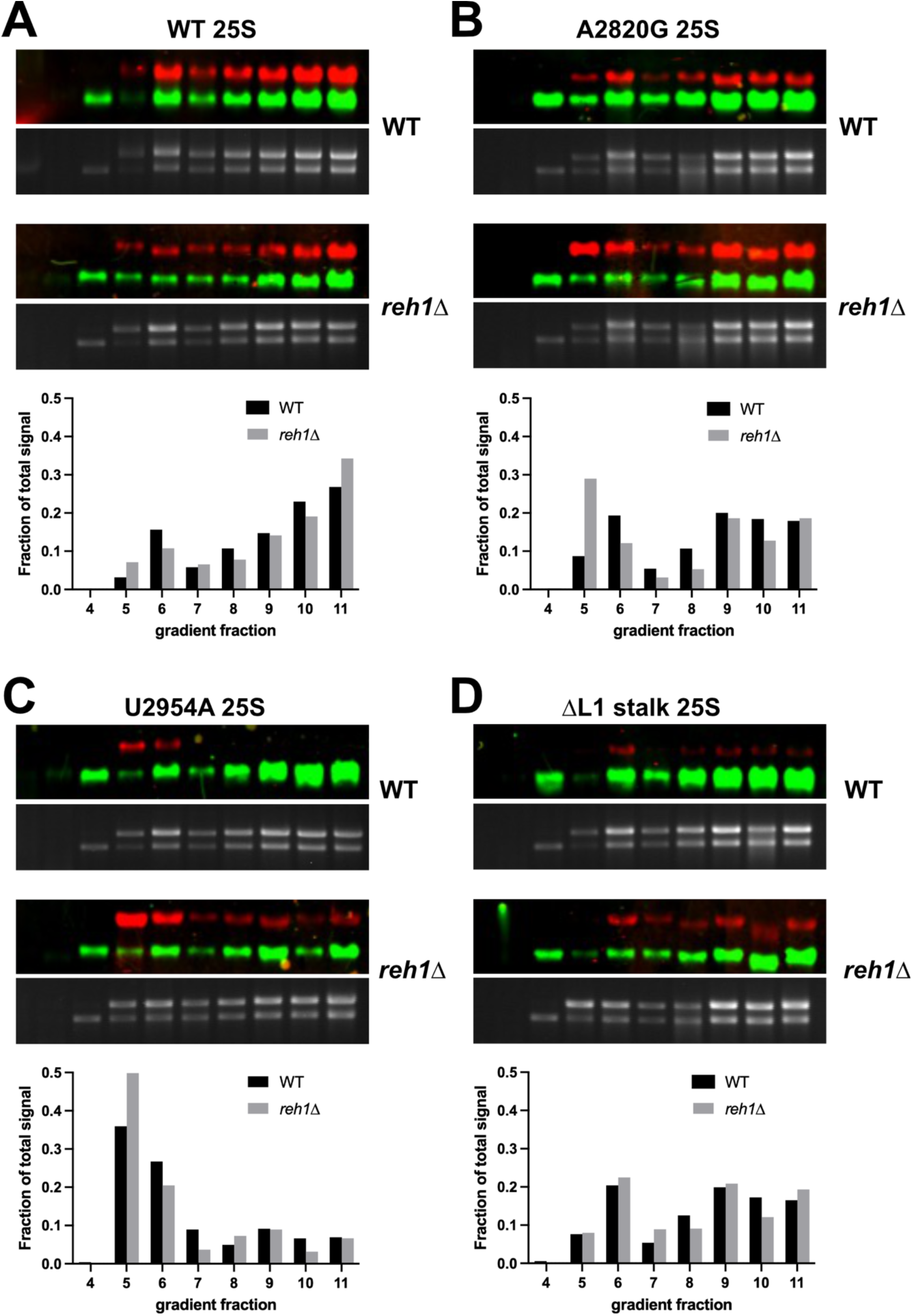
Sucrose gradient sedimentation analysis of mutant rRNA in WT and *reh1Δ* cells. Sucrose gradient density centrifugation was performed on extracts from wild-type or *reh1Δ* strains expressing 2µ plasmids with WT 18S rDNA and either WT, A2820G, U2954A, or L1Δ 25S rDNA. rRNA was extracted from each fraction and analyzed via northern blot. Total RNA from each fraction is shown by SYBR^TM^ safe. The proportion of signal contained in each fraction relative to the total signal is plotted. Analysis was based on single replicates.

## Discussion

Twenty years ago the Moore group showed that inactivating rRNA mutations in the catalytic center of the large ribosomal subunit or in the decoding center of the small subunit resulted in rapid degradation of the affected subunit[5]. We now know that defects in the small subunit lead ribosome stalling and collisions which are recognized by a dedicated machinery to clear the stalled ribosomes and associated mRNAs[28–30]. We also know that defective large subunits are removed by the 25S NRD pathway, which involves ubiquitination by the Rtt101 cullin E3 complex[6,31]. However, the ribosome-proximal factors that recognize defective 60S subunits to signal degradation and whether or not this pathway works during translation or at another step in ribosome metabolism have remained unanswered questions. Here, we show that in yeast, the zinc-finger protein Reh1 is required for 25S NRD and plays a role in identifying defecting subunits prior to them entering translation.

### *REH1* is required for 25S NRD

We previously identified Reh1 as the last biogenesis factor to associate with pre-60S subunits[32]. In fact, Reh1 can remain bound to 60S subunits until after they have engaged with a 40S subunit and initiated translation. At this stage, the C-terminus of Reh1 remains engaged in the polypeptide exit tunnel and its eviction appears to depend on the growing polypeptide chain[32]. Consequently, we previously proposed that Reh1 could recognize defective nascent 60S subunits by their inability to evict the C-terminus of Reh1 from the exit tunnel if they cannot carry out polypeptide synthesis[32].

To test if Reh1 targets defective ribosomes for degradation and whether this happens during biogenesis or translation, we utilized a defective ribosome model with a protein mutation in the P-site. During ribosome assembly in eukaryotic cells, the peptidyl transferase center is completed by the insertion of uL16, a late step in cytoplasmic maturation[17,18]. We have recently shown that 60S subunits that are assembled with mutant uL16 are targeted for degradation at the licensing step in biogenesis, in which the anti-association factor Tif6 is released[19]. In that work, we employed a mutant uL16 in which the P-site loop (aa 102-112) had been deleted, Because the P-site lop interacts with P-site ligands, including Sdo1, this mutant inserts into the ribosome but prevents the release of Tif6 which requires the P site ligand Sdo1[21,33]. A similar conclusion was reached for human ribosomes containing the analogous P site loop mutation in human uL16[24]. We also showed that mutant ribosomes that escape surveillance by Reh1 at this point become relatively stable as they enter translation[19]. Thus, the primary checkpoint for functionality of 60S subunits with mutant uL16 appears to be at the pre-60S licensing step and not during translation. Here, we have extended this result to show that Reh1 recognizes defects in or near the catalytic center, regardless of whether they are RNA or protein mutants.

We examined the expression and stability of 25S rRNA carrying two different mutations in the catalytic center of the ribosome, A2820G and U2954A (corresponding to the universally conserved A2451 and U2585 residues). We found that both mutations led to increased turnover of mutant 25S as previously reported[5,6]. However, we found that the magnitude of the impact of the rRNA mutations was less than previously reported when normalized to expression of 18S from the same vector. In addition, the two mutations impacted the stability of the mutant rRNAs to significantly different degrees, with the A2820G mutation having only a slightly reduced half-life compared to wild-type RNA and the U2954A having a severely reduced half-life. However, deletion of *REH1* fully stabilized both the A2820G and U2954A mutant rRNAs indicating that Reh1 is required for the degradation of these RNAs and suggesting that there is not another pathway in addition to Reh1 for degrading these defective 25S mutants. Thus, in yeast,

REH1 is essential for 25S NRD. Whether or not metazoan cells have pathways for recognizing and eliminating defective large subunits during translation remains an open question. It is possible that single-celled eukaryotes, such as yeast, have not evolved such pathways because cells containing a high flux of defective ribosomes in translation can be eliminated simply by growth competition.

As noted previously, the U2954A mutant does not appear to progress into polysomes, and arrests at the pre-60S stage, whereas a significant fraction of the A2820G mutant ribosomes progress into polysomes. We recently showed that ribosomes carrying a mutation in uL16 are relatively stabilized if they escape the biogenesis pathway and enter translation. We suggest a similar phenomenon is at play with the rRNA mutants; that they are surveilled by Reh1 during pre-60S maturation and not during translation. Indeed, we found that in the absence of *REH1* A2820G and U2954A mutant subunits accumulate as free-60S subunits, suggesting that these mutant ribosomes are eliminated prior to their entering translation. Previous work found that the ATPase Cdc48 was required for elimination of A2820G-containing ribosomes and that in the absence of Cdc48, A2820G subunits accumulate at 80S[16]. Those findings suggest that A2820G-containing subunits are recognized during translation and require Cdc48 for splitting the subunits prior to proteasomal degradation. We note that 25S carrying the A2820G mutation is fully stabilized in the absence of Reh1, indicating that Reh1 is required for its degradation. We previously suggested that the failure to evict Reh1 from the exit tunnel by the growing nascent chain could be a signal for a defective subunit[32]. Perhaps these subunits are split by Cdc48 and returned to the licensing step where Reh1 could recruit degradation machinery.

Surprisingly, not all defective 60S subunits appear to be subjected to surveillance and degraded. In particular, we found that subunits with a truncated L1 stalk display a half-life indistinguishable from wild-type ribosomes. The L1 stalk is a flexible appendage on the ribosome composed of an extended RNA element which provides the binding site for the ribosomal protein eL1[34]. The L1 stalk interacts with E site ligands during translation, including E site tRNAs and the translation elongation factor eIF5A as well as the nuclear export factor Nmd3[20,27,35]. Truncation of the L1 stalk RNA (ΔL1 stalk) removes the binding site for eL1 and is lethal in yeast[25]. These subunits are expressed at lower levels than wild-type, but they do engage with 40S subunits and enter polysomes. Their low expression is likely due to their inefficient export from the nucleus[25] but, the level of the ΔL1 stalk mutant rRNA is not affected by deletion of *REH1*. Deletion of *REH1* also does not alter the distribution of subunits in sucrose gradients. The ability of ΔL1 stalk mutant 60S to escape surveillance and engage in translation is reminiscent of findings for 60S subunits in yeast depleted of Las1, which is required for removal of internal transcribed spacer 2 (ITS 2) from the 27SB pre-rRNA[36–38]. Las1 depletion results in 60S subunits that retain ITS2 and associated factors, forming a “foot”-like appendage on the subunit[39]. These subunits join 40S subunits and are catalytically active. They also recruit the ribosome-associated quality control factors Rqc2 and Ltn1, involved in the removal of nascent peptides from stalled ribosomes[40–42]. However, direct evidence that these aberrant “foot”-containing 60S are selectively degraded is lacking. Based on our work reported here, we conclude that surveillance pathways for defective 60S subunits in yeast have restricted substrate recognition. Mutations in or near the catalytic center appear to be handled by an Reh1-dependent 25S NRD pathway but ΔL1 stalk mutant ribosomes appear immune to clearance. It remains to be tested if other classes of defective 60S subunits, such as the “foot”-containing ribosomes from *LAS1* mutants are recognized by Reh1 or by other unidentified mechanisms.

### How does Reh1 detect defects in the catalytic center?

The N-terminus of Reh1 binds to the surface of Tif6 on the joining face of the pre-60S subunit[19]. The remainder of the protein wraps around the subunit and the C-terminus enters the exit tunnel, approaching the PTC[17,19,43]. One could imagine that the C-terminus of Reh1 detects subtle allosteric changes induced by mutations in the PTC. However, we think a more likely scenario is that Reh1 communicates with the PTC through Sdo1, which binds in the P-site, and Efl1 which simultaneously spans Sdo1 and Tif6[44]. We propose that mutations in the PTC perturb the binding of Sdo1 in such a way that it is unable to activate the GTPase Efl1 to evict Tif6. Thus, the persistence of Reh1 on the ribosome post-licensing may be the signal for subunit degradation.

### How does Reh1 communicate with downstream degradation machinery?

Elegant work from the Kitabatake and Ohno labs has shown that degradation of 25S NRD substrates requires the cullin-E3 ligase Rtt101, which acts with Mms1, Hrt1 and Cdc34 to ubiquitinate components of the 60S subunit to target them for proteasomal degradation[6,31]. The Rtt101-Mms1 complex works in both DNA repair and 25S NRD pathways, however different substrate-specific adapters are recruited for each pathway. Mms22 is the adapter for substrate recognition in DNA repair, whereas Crt10 is the adapter in 25S NRD[31]. It is not currently known what substrate Crt10 recognizes that leads to 25S NRD, however the role of Reh1 as a quality control factor required for degradation of aberrant 25S rRNA, shown here, implicates it as a prime candidate.

Additionally, *REH1* is expressed distinctly from canonical ribosome biogenesis factors, instead being co-expressed with factors involved in proteostasis including the proteasome[19]. Finally, the proximity of Reh1 to the exit tunnel is provocative. The exterior of the exit tunnel is a site known to be associated with quality control factors, such the Ribosome-Associated Quality Control factor Ltn1 recognizes nascent polypeptides on stalled ribosomes[45]. Whether or not Reh1 directly recruits the RTT101 complex, what proteins are ubiquitinated to initiate 25S NRD, and whether they are ribosomal proteins or associated assembly factors such as Reh1 itself, are questions remaining to be resolved.

### Does ZNF622, the human homolog of Rei1/Reh1, play a role in quality control and 25S NRD?

S. cerevisiae, and a subset of hemiascomycetes contain Reh1 and a paralog Rei1 which resulted from an ancient genome duplication[46]. With the exception of some plants, which have two copies of Rei1 resulting from an independent gene duplication[47], most eukaryotes contain a single gene, which is ZNF622 in humans.[47] In yeast, Rei1 and Reh1 have both separate and redundant functions. Rei1 but not Reh1, recruits Ssa and its associated J-protein Jjj1 to promote the removal of the export factor Arx1[48–50]. Rei1 and Reh1 together share a redundant role in the loading of the ribosomal protein eL24[51] while only Reh1 acts as a quality control factor[19]. It is not currently known if ZNF622 plays a quality control role in human cells. However, it is intriguing that ZNF622 removal from pre-60S is dependent on ubiquitination by the E3 ligase HECT D1[52]. Knockdown of HECT D1, leads to accumulation of ZNF622 and eIF6 on 60S subunits, suggesting that ZNF622 remains associated with pre-60S through SBDS-EFL1-dependent eviction of eIF6[52]. This places ZNF622 at a point in the 60S maturation pathway where it could serve as a surveillance factor akin to Reh1. It is also probable that 25S NRD in higher eukaryotes is more complex than in yeast.

Indeed, a recent screen for surveillance factors identified ZNF574 as a quality control factor that eliminates defective uL16[24]. ZNF574 does not have a clear counterpart in yeast and whether or not ZNF574 is required for surveillance of 25S NRD substrates remains an open question.

## Acknowledgements

This work was funded by NIH grant GM127127 to A.W.J. The authors than P. Huang and E. Kypri for help with assembling vectors and F. LaRiviere for providing the pJV series of vectors used here.

## Materials and Methods

### Yeast strains, plasmids and cell growth

Yeast strains, plasmids and oligonucleotides used in the present study are listed in Tables 1-3, respectively. AJY4949 and AJY4950 were derived by sporulating and dissecting the respective heterozygous deletion mutants (Research Genetics). AJY1185 was made by swapping pAJ724 into JD1111. AJY1661 was made by swapping pAJ1181 into AJY1185. pAJ1181 was made by inserting the sequence GAAGAGCCATTGCACTCCGGTTCTTCTGCAG between nucleotides 137 and 138 of 25S rDNA in pJD211.LEU. pAJ5650 and pAJ5659 were made by moving the Nde1 Sph1 fragment containing tagged 18S from pWL109 into pAJ3605 and pAJ1181, respectively. pAJ4948, pAJ5658, pAJ5660, pAJ5664, and pAJ4988 were made by introducing the indicated point mutations into pAJ5659. pAJ5651, pAJ5662 and pAJ5663 were made by PCR amplification of the 25S region from pAJ1181, pAJ1963 and pAJ3605, respectively, using oligonucleotides AJO664 and AJO4414 followed by Sph1, SalI digestion and cloning into the same sites of pWL109. pAJ5764 was made by moving the SacI-MluI fragment from pAJ5651 into the same sites of pAJ5666. All rDNA vectors were sequenced in their entirety. pAJ1963 was made by site directed mutagenesis in pJD211.LEU. All cell cultures were grown in synthetic dropout media[53] supplemented with 2% glucose unless noted otherwise.

**Table 1:**

| Strain | Genotype | Source |
| --- | --- | --- |
| AJY1185 | *MATa ade2-1 ura3-1 leu2-3 his3-11 trp1 can1-100 rdn1ΔΔ::HIS3/pJD209.TRP*, pAJ724 | [54] |
| AJY1661 | *MATa ade2-1 ura3-1 leu2-3 his3-11 trp1 can1-100 rdnΔΔ::HIS3/pJD209.TRP*, pAJ1181 | This study |
| AJY4618 | *MATa reh1Δ::Natr his3Δ1 leu2Δ0 ura3Δ0* | [19] |
| AJY4941 | *MATa ade2-1 ura3-1 leu2-3 his3-11 trp1 can1-100 rdnΔΔ::HIS3/pJD209.TRP*, pAJ5659 | This study |
| BY4741 | *MATa his3Δ1 leu2Δ0 ura3Δ0 met15Δ0* |  |
| JD1111 | *MATa ade2-1 ura3-1 leu2-3 his3-11 trp1 can1-100 rdn1ΔΔ::HIS3/pJD209*, pJD211.LEU | [54] |

**Table 2:**

| Plasmid | Features | Source |
| --- | --- | --- |
| pAJ724 | 25S-U1A URA3 2 micron Amp | [55] |
| pAJ1181 | 25S-U1A LEU2 2 micron Amp | [25] |
| pAJ1963 | 25S-U1A-A2820G LEU2 2 micron Amp | This study |
| pAJ3605 | 25S-U1A-ΔL1 stalk LEU2 2 micron Amp | [25] |
| pAJ4988 | 25S-U1A 18S-tag-A1775C LEU2 2 micron Amp | This study |
| pAJ4993 | 25S-oligo(pJV12) 18S-oligo(pJV12) LEU2 2 micron Amp | This study |
| pAJ4994 | 25S-oligo(pAJ5669) 18S-oligo (pJV12) LEU2 2 micron Amp | This study |
| pAJ5651 | GAL7:rDNA 18S-oligo and 25S-U1A URA3 2 micron Ampr | This study |
| pAJ5652 | GAL7:rDNA 18S-oligo and 25S-U1A-A2820G URA3 2 micron Ampr | This study |
| pAJ5653 | GAL7:rDNA 18S-oligo and 25S-U1A-ΔL1 stalk 2 micron URA3 Ampr | This study |
| pAJ5658 | 25S-U1A-A2820G 18S-tag LEU2 2 micron Amp | This study |
| pAJ5659 | 25S-U1A 18S-tag LEU2 2 micron Amp | This study |
| pAJ5660 | 25S-U1A-ΔL1 stalk 18S-tag LEU2 2 micron Amp | This study |
| pAJ5661 | 25S-U1A 18S-tag LEU2 CEN Amp | This study |
| pAJ5662 | 25S-U1A-A2820G 18S-tag LEU2 CEN Amp | This study |
| pAJ5663 | 25S-U1A-ΔL1 stalk LEU2 CEN Amp | This study |
| pAJ5665 | 25S-U1A-U2954A 18S-tag LEU2 CEN Amp | This study |
| pAJ5664 | 25S-U1A-U2954A 18S-tag LEU2 2 micron Amp | This study |
| pAJ5764 | GAL7:rDNA 18S-oligo 25S-U1A-U2954A URA3 2 micron Ampr | This study |
| pJD209.TRP | 5S rDNA TRP1 Amp | [54] |
| pJD211.LEU | 35S rDNA LEU2 Amp | [54] |
| pJV12 | pPGK1-rDNA LEU2 2 micron Amp | [5] |
| pJV12(A2451G) | pPGK1-rDNA-A2820G LEU2 2 micron Amp | [5] |
| pJV12(U2585A) | pPGK1-rDNA-U2954A LEU2 2 micron Amp | [5] |
| pWL109 | GAL7:rDNA (18S-tag) URA3 2 micron Amp | [56] |

**Table 3.**
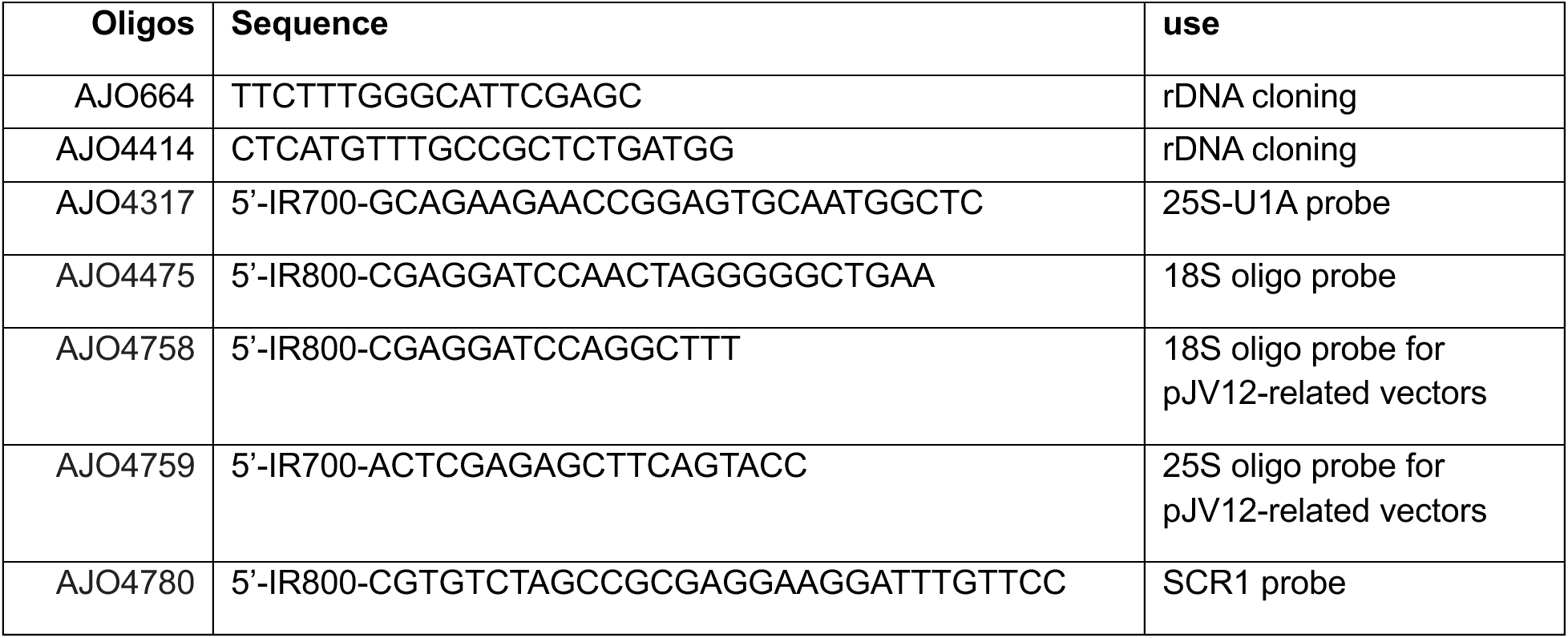

### Steady State RNA Analysis

All analyses were done in triplicate from independent yeast colonies. Fresh overnight cultures were diluted to OD600 of 0.05 (or 0.1 for slow growing stains) in 10 mL of selective medium, as indicated, and grown with shaking at 30°C to OD600 of 0.3-0.5. Cells were collected by centrifugation and stored at −80°C until use. RNA was extracted by the hot phenol method using Phaselock (5-Prime) as described[57]. RNAs were resuspended in nuclease-free water and quantified by absorbance at 260 nm.

### RNA time course

Cultures of the indicated strains and vectors were grown to OD600 of 0.3 in synthetic dropout medium containing 1% galactose. At time zero, cells were pelleted and resuspended in synthetic dropout media containing 2% glucose and 10 ml samples were harvested at 0, 30, 60 and 120 minutes after resuspension.. The OD600 was measured as samples were collected. Samples were flash frozen and store at −80°C prior to extraction of RNA and analysis as described for Steady State RNA analysis

### Northern Blotting

Either 5ug total RNA (for strains containing 2 micron vectors) or 10µg total RNA (for strains containing CEN vectors) was loaded per sample except in the case of sucrose density gradient analysis for which constant fraction volumes were loaded. RNAs were electrophoresed through 1% agarose-formaldehyde gels in MOPS buffer. RNAs were transferred to Brightstar^TM^ nylon membrane by capillary action and crosslinked in a UV Stratlinker 1800 (Stratagene) using the auto crosslink setting. Membranes were probed with 10 pmol of probe/10 mL of hybridization solution. Northern blots were imaged on an Odyssey CLx imager in the 700 and 800 nm channels. Images were quantified in Image Studio (Li-Cor) and data were compiled in Microsoft Excel and analyzed in Prism (GraphPad). Significance was determined using multiple unpaired t-tests. Total RNAs were analyzed by electrophoresis through 1% agarose gels in TAE buffer containing Sybersafe^TM^ and imaged.

### Sucrose density gradient sedimentation

Fresh overnight cultures were diluted to OD600 of 0.05 in 50 mL of selective medium, as indicated, and grown with shaking at 30°C to OD600 of 0.3-0.5. Cycloheximide (VWR life sciences) was added to a final concentration of 100 μg/mL, and cultures were incubated with shaking for an additional 10 min at 30°C prior to harvesting. Cell pellets were immediately washed and resuspended in cold 150 μL of lysis buffer containing 20 mM Tris·HCl (pH 7.5), 100 mM NaCl, 30 mM MgCl_2_100 μg/mL CHX, 200 μg/mL heparin, 5 mM β-mercaptoethanol, 1 mM PMSF, and 1 μM each of leupeptin and pepstatin). Cell extracts were prepared by vigorous agitation with glass beads three times for 25 seconds, with intervals on ice. Supernatant was removed and beads were washed with 100 uL buffer. Supernatant and wash were combined and clarified by centrifugation at 18,000 × g for 10 min at 4°C. Clarified extract corresponding to 5 A_260_ units was layered onto 10–60% (w/v) sucrose gradients prepared in TMN buffer (50 mM Tris-acetate, pH 7.0, 50 mM NH_4_Cl, 12 mM MgCl_2_) and centrifuged for 70 min at 50,000 rpm at 4°C in a Beckman SW55 rotor. Gradients were fractionated using a Biocomp Piston Gradient Fractionator fitted with a Triax flow cell, at 3 mm/s. Gradients were monitored at 260 nm and RNA was precipitated as described[58]. Pellets were resuspended in 25uL of nuclease-free water. 25 uL of 2X RNA loading dye (NEB) was added and samples were heated at 65°C for 8 min. 10 uL of each sample was loaded per lane.

---

**Supplemental Figure S1.**
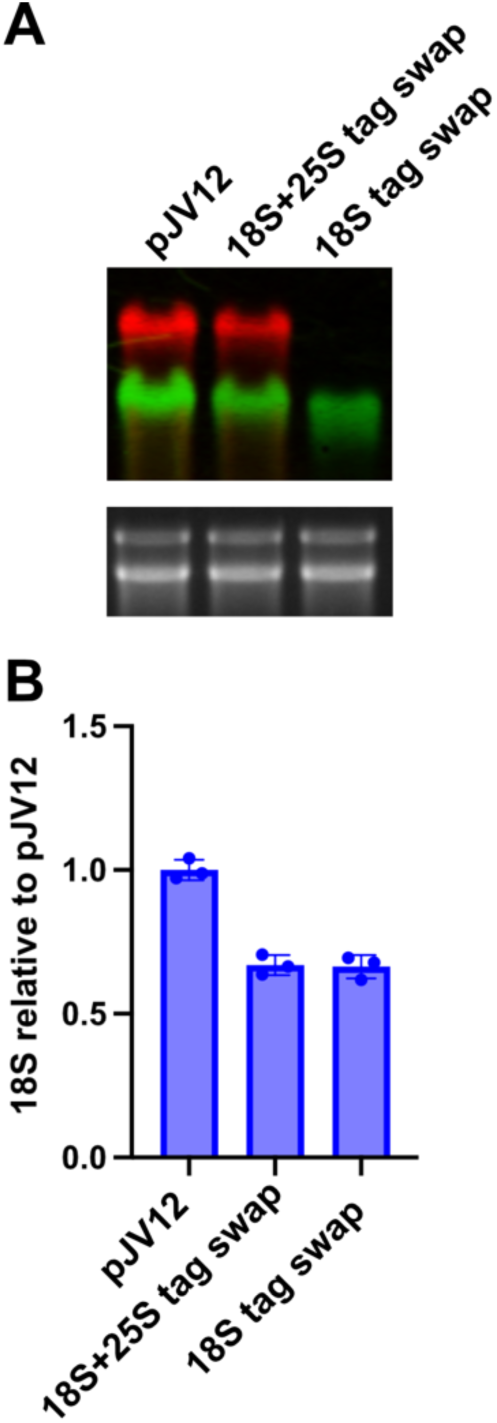
Comparison of expression of the pJV12 series PGK1 vectors and the Pol I vectors generated for this work. For direct comparison, both 25S and 18S oligo tags from pJVJ12 were ported into the RNA Pol I vector (pAJ4993, 18S + 25S tag swap) or only the 18S tag was swapped (pAJ4994, 18S tag swap). **A.** RNA was prepared from WT cells expressing the indicated vectors and analyzed by northern blotting with oligonucleotide probes AJO4758 and AJO4759. Total RNA was stained by SYBR safe (lower panel). The absence of 25S signal in the 18S tag swap reflects the fact that the probe for pJV12 does not recognize the tag in pAJ4994. **B.** Quantification of 18S levels relative to pJV12. Quantitation was from three experimental replicates.

**Supplemental Figure S2.**
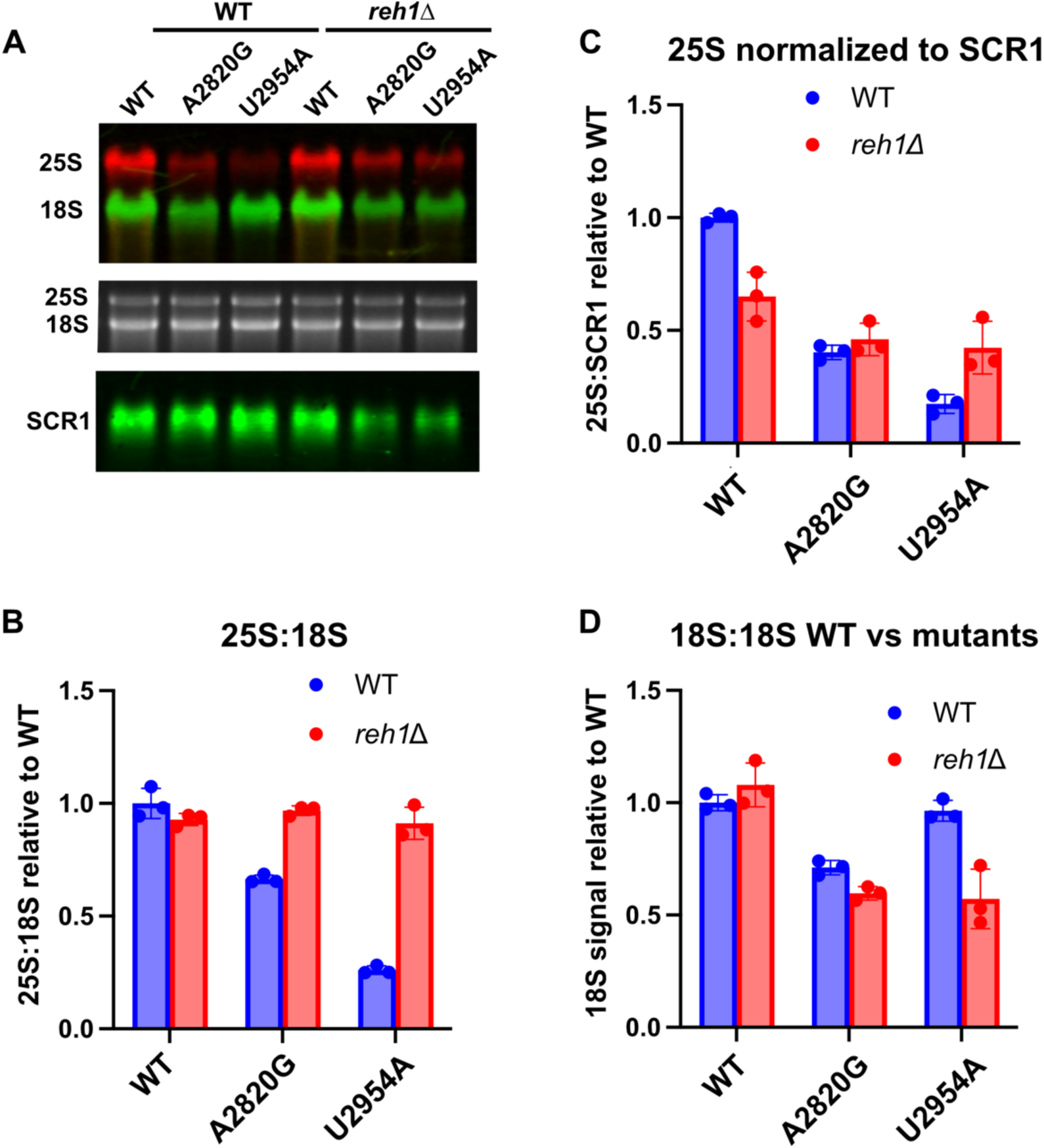
WT and mutant rRNA were expressed in WT and *reh1*Δ cells under control of the *PGK1* promoter from vectors described previously[5]. **A.** Representative northern blot using probes AJO4758 and AJO4759 for 18S and 25S rRNA, respectively, and AJO4780 for *SCR1* as a loading control. Equal amounts of total RNA were loaded per lane. Middle panel shows total RNA stained with SYBR^TM^ safe. **B.** Quantification of 25S:18S ratios normalized to WT rRNA in WT cells. Data are from three experimental replicates. **C.** Quantitation of 25S levels when normalized to *SCR1*. **D.** Comparison of 18S levels normalized to 18S from WT rRNA in WT cells. Quantitation was from three experimental replicates.

**Supplemental Figure S3.**
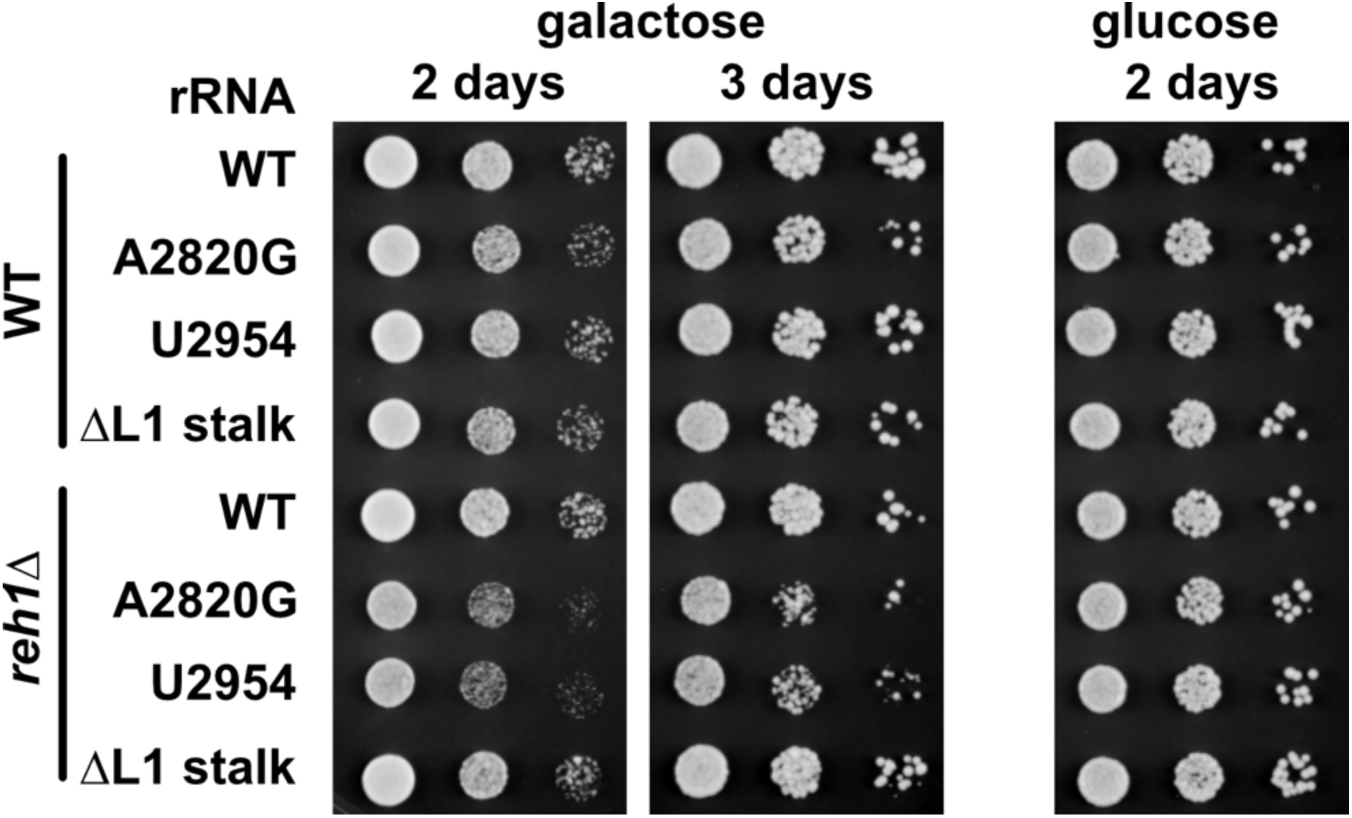
Growth assay of WT (BY4741) and *reh1Δ* (AJY4618) cells expressing galactose-inducible WT or mutant rDNAs. 10-fold serial dilutions of cells were spotted onto galactose- or glucose-containing SD Leu- medium and plates were imaged after the indicated times at 30°C.

**Supplemental Figure S4.**
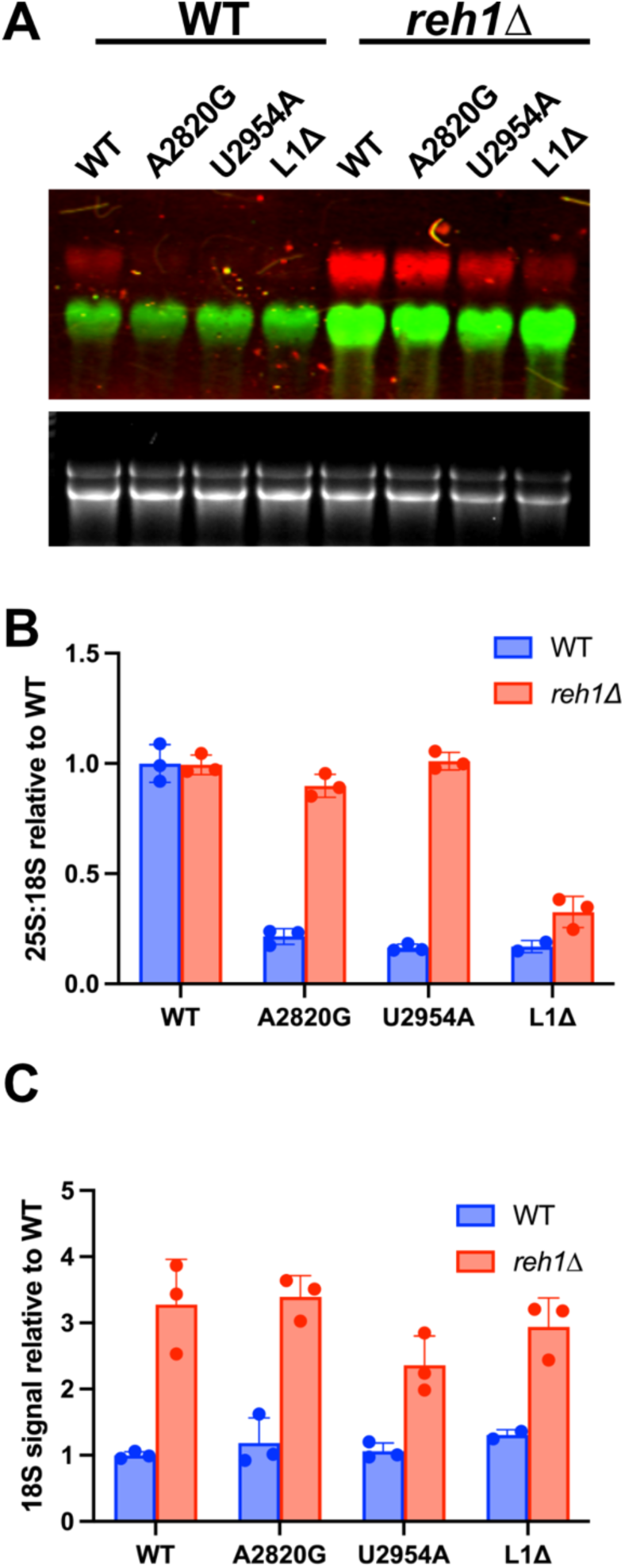
Expression from low copy vectors results in lower steady state levels of mutant rRNAs. **A.** WT or mutant rRNAs were expressed in WT or *reh1Δ* cells under control of the native RNA Pol I promoter but from low copy centromeric vectors. RNA was prepared and analyzed by northern blotting as described in the legend to Figure 1. Equal amounts of total RNA were loaded in each lane. Upper panel, two color fluorescent northern blot. Lower panel, SYBR safe-stained RNA. **B.** Quantification of the ratio of 25S to 18S, normalized to the 25S to 18S ratio of WT rRNA in WT cells. RNAs were prepared from three experimental replicates. **C.** Quantification of the ratio of 18S levels of WT and mutant rRNA in WT and *reh1Δ* mutant cells compared to WT rRNA in WT cells. Quantitation was from three experimental replicates.

**Supplemental Figure S5.**
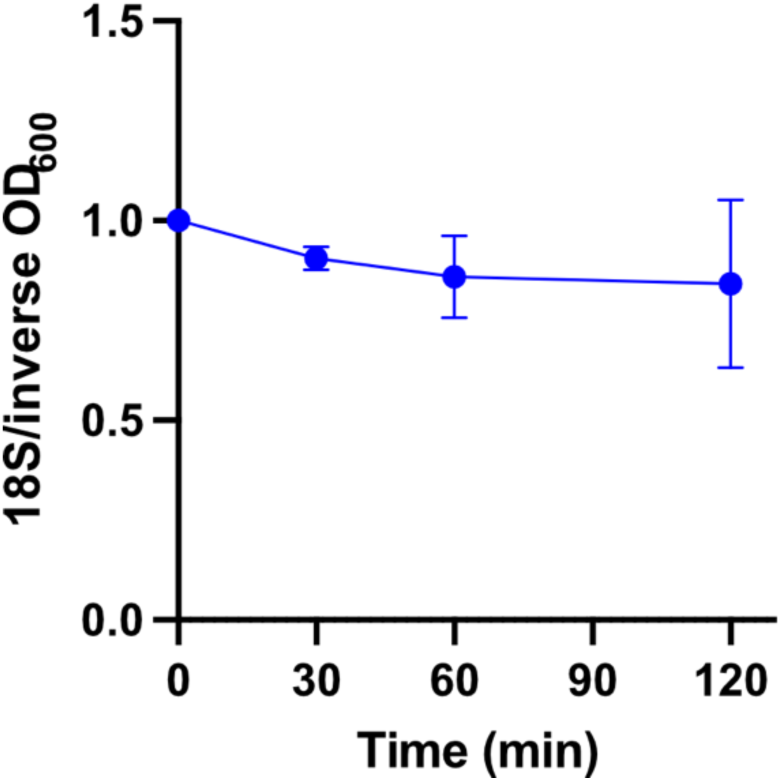
Quantification of WT 18S levels in WT cells from Figure 3A plotted as 18S signal over the inverse of cell density over time. Quantitation was from three experimental replicates.

